# Transcriptome network analysis link perinatal *Staphylococcus epidermidis* infection to microglia reprogramming in the immature hippocampus

**DOI:** 10.1101/2022.07.04.498695

**Authors:** Giacomo Gravina, Maryam Ardalan, Tetyana Chumak, Halfdan Rydbeck, Xiaoyang Wang, Carl Joakim Ek, Carina Mallard

## Abstract

*Staphylococcus epidermidis* (*S. epidermidis*) is the most common nosocomial pathogen in preterm infants and associated with increased risk of cognitive delay, however, underlying mechanisms are unknown. We employed morphological, transcriptomic and physiological methods to extensively characterize microglia in the immature hippocampus following *S. epidermidis* infection. 3D morphological analysis revealed activation of microglia after *S. epidermidis*. Differential expression combined with network analysis identified NOD-receptor signalling and trans-endothelial leukocyte trafficking as major mechanisms in microglia. In support, active caspase-1 was increased in the hippocampus and using the LysM-eGFP knock-in transgenic mouse, we demonstrate infiltration of leukocytes to the brain together with disruption of the blood-brain barrier. Our findings identify activation of microglia inflammasome as a major mechanism underlying neuroinflammation following infection. The results demonstrate that neonatal *S. epidermidis* infection share analogies with S. aureus and neurological diseases, suggesting a previously unrecognized important role in neurodevelopmental disorders in preterm born children.

## Introduction

*S. epidermidis*, a coagulase-negative staphylococci (CoNS), is the most common nosocomial pathogen in neonatal intensive care units, responsible for up to 50% of all cases of late-onset sepsis in neonates (Dong and Speer, 2014). The incidence of CoNS infections is inversely associated with neonatal maturity, being highest in very low birth weight infants (Boghossian et al., 2013) and evidence shows that CoNS sepsis in preterm infants is associated with increased risk for cognitive delay (Alshaikh et al., 2014). Neonatal bloodstream infections can trigger neuroinflammation and are believed to contribute to neurological outcome in preterm infants (Hagberg et al., 2015, Strunk et al., 2014). Systemic *S. epidermidis* infection induces increased white blood cell count in the CSF and chemokine levels in the brain as well as reduced volume of grey and white matter in neonatal mice (Bi et al., 2015) and proinflammatory proteome changes in plasma and CSF in newborn preterm pigs (Muk et al., 2019). We later also demonstrated that an ongoing *S. epidermidis* infection sensitizes the neonatal brain to hypoxic-ischemic (HI) injury (Lai et al., 2020, Gravina et al., 2020).

While microglia is involved in several crucial steps of normal brain development, such as synapse function, plasticity, and circuit formation (Ardalan et al., 2019), in neurological conditions microglia transcriptional program can shift towards a proinflammatory injurious state (Cserep et al., 2021). It has been suggested that neuroinflammation is an important factor in *S. epidermidis*-induced brain injury, however, the role of microglia in these processes remain unclear (Joubert et al., 2022). Therefore, we sought to characterize the cellular and molecular phenotype of microglia in the neonatal mouse hippocampus following systemic *S. epidermidis* infection. As microglia show regional heterogeneity (Tan et al., 2020), we specifically focused on the hippocampus because hippocampal development is impaired in very preterm infants and associated with memory and cognitive deficits (Beauchamp et al., 2008, Strahle et al., 2019), and it retains neurogenesis in the adult brain (Fares et al., 2019). Thus, injury to the hippocampus in the perinatal period might have long-term effects on the brain. 3D analysis of microglia morphology showed activation following *S. epidermidis* infection. Employing global transcriptome analysis of microglia, we identified activation of NOD-signalling and the inflammasome following *S. epidermidis* infection. Network analysis in combination with differential gene expression analysis revealed overrepresentation of biological processes involved in regulation of leukocyte activation. In support, physiological in vivo experiments showed increased permeability of the blood-brain barrier (BBB) and infiltration of myeloid cells, suggesting importance of microglia-leukocyte communication in the activation of the microglia inflammasome following neonatal *S. epidermidis* infection.

## Results

### 1. S. epidermidis infection induces morphological changes in hippocampal microglia

As perinatal infection and inflammation can affect brain injury in a sex-dependent manner (Ardalan et al., 2019), we evaluated microglia morphology transformation associated with activation following peripheral *S. epidermidis* infection in the hippocampus in male and female pups separately. In physiological conditions, resting microglia exhibit ramified cell morphology with long thin processes extending from their spherical soma. Upon activation, microglia arborisations are fewer and shortened resulting in a rounded amoeboid-like appearance (Cserep et al., 2021). Three-D analysis showed that the soma volume of Iba1 positive microglia in the CA1 stratum radiatum (CA1SR) hippocampal subregion was significantly increased after *S. epidermidis* infection (Fig 1A&B). Specifically, a larger microglia soma was observed in infected male mice compared to saline male mice (p<0.0001, Fig 1B). The total length and the number of microglia branches were profoundly altered by *S. epidermidis* infection (Fig 1 C-E). Comparison between saline and infected mice showed a decrease of microglia branches in both males and females following *S. epidermidis* infection (p<0.0001 and p=0.002, respectively). Consistent with cell soma volume, changes in microglia soma sphericity were identified in *S. epidermidis* male mice but not in female animals (Fig 1F&H). To further evaluate the morphological alteration of microglia following *S. epidermidis* infection, we compared the branching complexity pattern of microglia. The results showed a significant reduction in the microglia arborisation in *S. epidermidis* infected mice compared to the saline group. Significantly lower number of branching intersections from the cell soma were observed in male mice at distances of 15-60 μm from soma while in females it was observed at the distance of 10-30 μm from soma (Fig 1G). Of note, differences in baseline levels were observed between male and female in soma volume, sphericity and total length of branches (p=0.001, p=0.008 and p= 0.009, respectively) (Fig 1B,D,F) A similar pattern of microglia activation as observed in the CA1SR was found in the molecular layer of dentate gyrus (MDG) hippocampal subregion (Fig S1).

**Figure 1.**
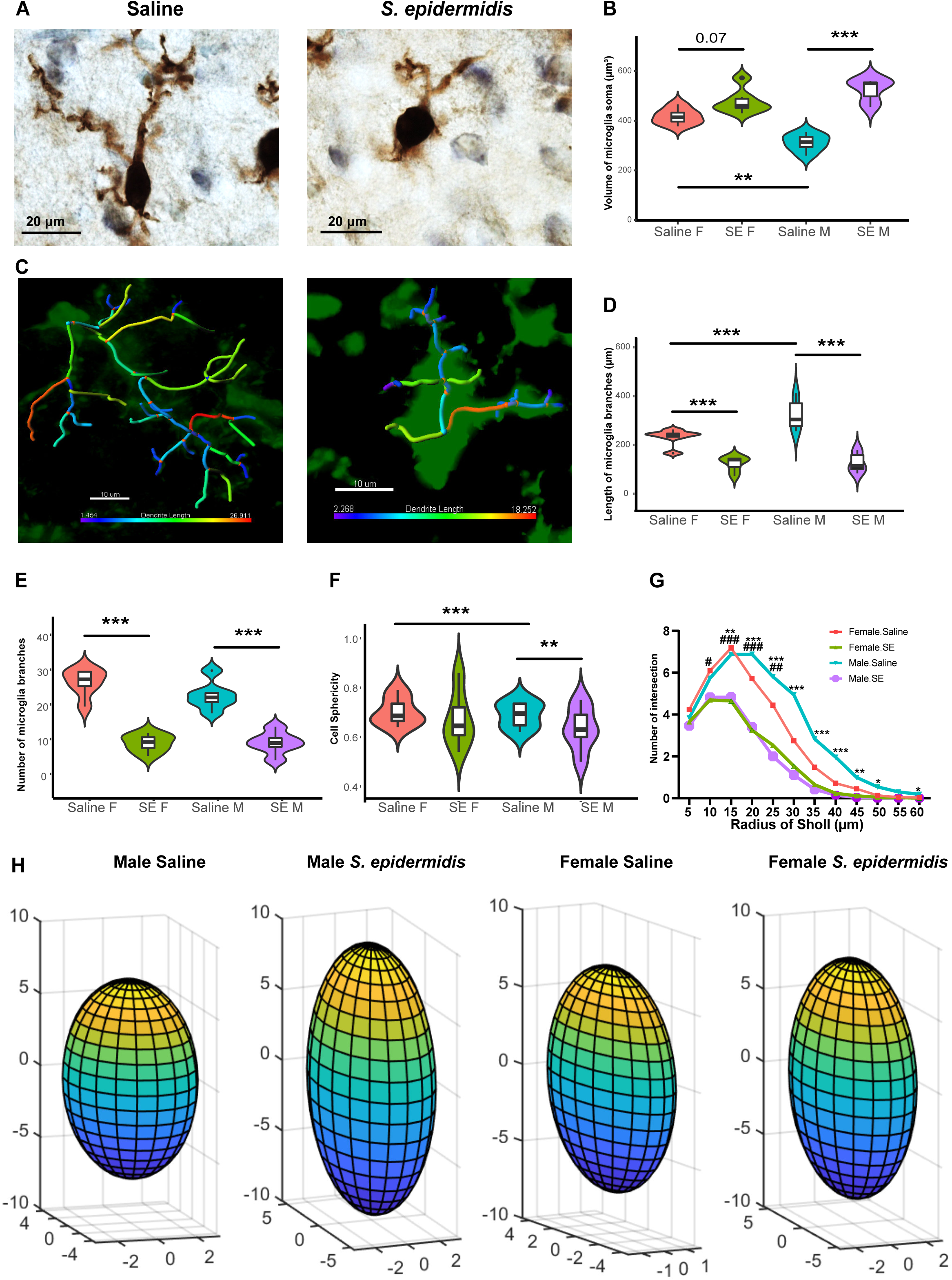
Activation of hippocampal microglia cells following S. epidermidis infection in neonatal mice. Representative images of microglia cell soma (100× oil immersion objective lens) (A) and 3D reconstructed microglia (C) in sections stained with Iba-1 of saline (left) and *S. epidermidis* (right) mice. Microglia volume (B), and branches length (D), number (E) and soma sphericity (F) was quantified in PND5 mice (n=6 sex/treatment). (G) Sholl analysis is indicating the complexity of microglia branching ramifications in the CA1SR hippocampal subregion. Significant differences between *S. epidermidis* and saline are indicated in * for male mice and # for female mice (H) 3D ellipsoid plots of microglia soma sphericity were generated for each group. Data are presented as median, 10–90th percentile, kernel probability density (violin). Statistical comparison between the *S. epidermidis* and saline groups for each parameter was performed using Two-way ANOVA with Tukey’s multiple comparison post-hoc test. *p<0.05; **p<0.01; ***p<0.001

### 2. S. epidermidis induces a distinct transcriptional signature in microglia

As morphological analysis indicated activation of microglia following *S. epidermidis* infection, we isolated microglia from the hippocampus of PND5 mice and performed RNAseq to characterize associated molecular changes (Fig 2A). Saline and *S. epidermidis* animal groups were clearly identified by hierarchical clustering of microglia transcripts (Fig 2B) and principal component analysis of the gene expression profiles (Fig 2C). The separation of gene expression profiles between the treatment groups consisted of 769 differentially expressed genes (DEGs). Out of these, 344 were upregulated and 425 downregulated in *S. epidermidis* infected pups (Fig 2D) with the top 10 up and down-regulated genes highlighted in a heatmap (Fig 2E). There were no significant sex-dependent effects on differentially expressed genes (Fig S2).

**Figure 2.**
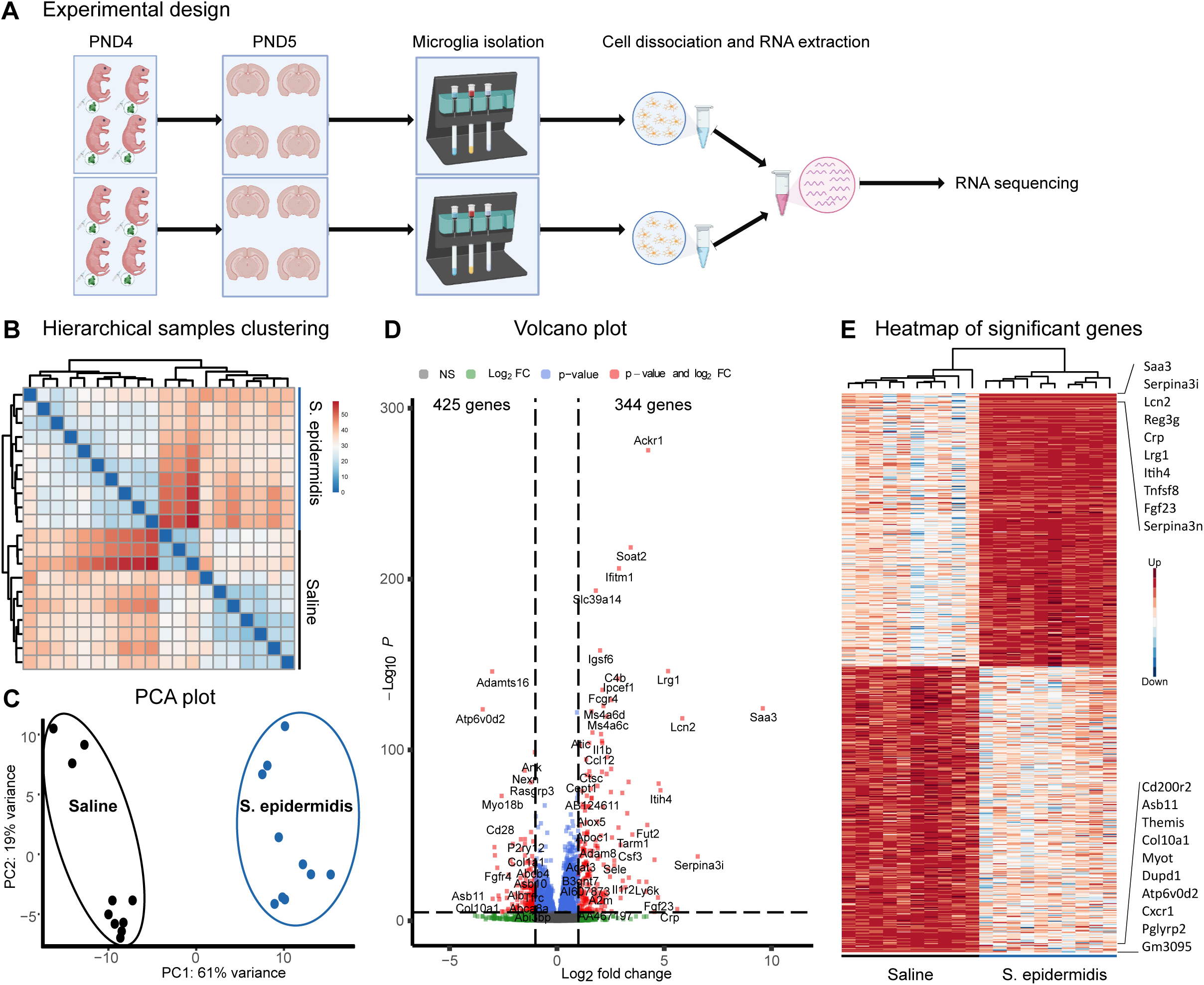
Transcriptomic changes of hippocampal microglia following S. epidermidis infection. An overview of microglia isolation protocol and RNA sequencing performed (A). Five samples per sex/group were used in the analysis. Each sample contained hippocampi deriving from 8 mice (same sex/treatment), thus a total of 160 mice were used for the sequencing. Hierarchical clustering of hippocampal microglia RNA-seq analysis using Euclidean distance reveal a separation of saline and *S. epidermidis* (saline n = 10, *S. epidermidis* n = 10) (B). Principal component analysis (PCA) also shows clear separation of microglia transcripts of saline and *S. epidermidis* at 24 h post infection (saline n = 10, *S. epidermidis* n = 10) (C). Volcano plot showing the spread of 21832 differentially expressed genes, 325 upregulated and 349 downregulated. Dotted lines represent the cut-values (padj < 0.05, FC > 2) (D). Heatmap of differentially expressed genes (E). The top 10 up and downregulated genes are enlarged and listed in the heatmap.

### 3. S. Epidermidis induces neuroinflammatory responses in the hippocampus

To gain insights into the molecular mechanisms and pathways associated with differential gene expression, we performed gene ontology (GO) enrichment analysis. We analysed up- and down-regulated genes separately, as it has been shown to be more powerful than combined analysis of all DEGs together (Hong et al., 2014). Analysis of biological processes of the upregulated genes revealed multiple immune related signatures, such as Defense response (GO:0006952), Immune system process (GO:0002376), Inflammatory response (GO:0006954), Defense response to other organism (GO:0098542) (Fig. 3A), while analysis of the downregulated genes revealed alterations in developmental processes, including System development (GO:0048731) and Anatomical structure development (GO:0048856) (Fig 3B). To identify potential signalling pathways associated with *S. epidermidis* infection, we annotated the regulated genes by KEGG pathway analysis. Intriguingly, we found that it was mainly the upregulated genes that were associated with KEGG pathways, such as Cytokine-cytokine receptor interaction, Staphylococcus aureus infection, Complement and coagulation cascades, IL-17 signalling pathway, NOD-like receptor signalling pathway (Fig 3C). Of note, S. aureus infection (more virulent strain of the Staphylococcus family) was one of the most enriched pathways, suggesting similarity in the inflammatory response between these bacterial infections. To gain further mechanistic insight into the upregulated and downregulated network, gene-set enrichment map analysis was performed (Merico et al., 2010). We found significant enrichment in multiple categories, particularly in up-regulated genes, including regulation of leukocytes and lymphocytes, as well as changes in ion homeostasis (Fig S3). An interactome network identified *Sell, Fcgr3, Fcgr4, Il1b, Cd14, Fcgr2b, C3, Crp, Apob, Csf3* as hub genes in the upregulated network (Fig. 3D & Table S1), while *Il7r, Cd28, Col1a1, Itgax, Col3a1, Itga2, Igf1, Il13, Acta2, Alb* as hub genes in the down-regulated network (Fig. 3D & Table S2).

**Figure 3.**
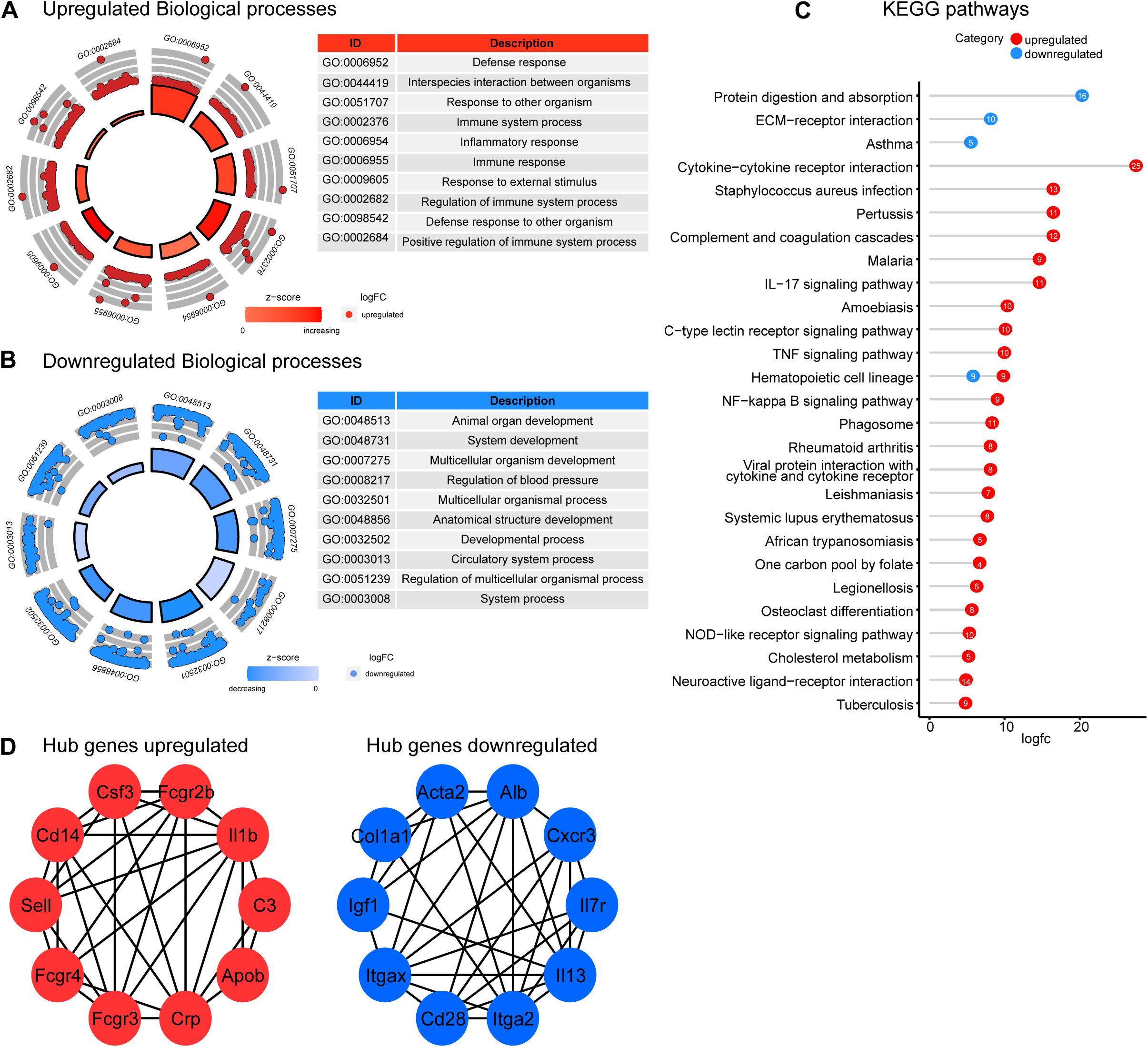
Functional enrichment analysis of differential expressed microglia genes. GO Plot representation of the top 10 significant GO terms in the upregulated (A) and down-regulated (B) network, with GO term descriptions in the table on the side. The outer circle represents how many genes are included in the GO term. The height of the bar indicates the negative log10 adjusted *P*-value. Significantly enriched KEGG pathways of the upregulated (red) and downregulated (blue) genes (C). The number inside the dot represents the number of genes enriched in the KEGG pathway. The top 10 high-degree genes screened from the PPI network by using MNC algorithm (D).

### 4. Transcriptional profile of microglia cells revealed by network analysis

To capture the full extent of gene expression profiles of microglia cells after *S. epidermidis* infection from a systems perspective, we performed weighted gene co-expression network analysis (WGCNA). WGCNA is a useful method to link tightly co-expressed gene modules to phenotypic traits (e.g. sex and infection) (Langfelder and Horvath, 2008). The full dataset of 20 samples (n=5/sex/treatment) was used for WGCNA analysis. No outliers were identified; therefore, all samples were included in the analysis. In total, 28 modules were identified and most of the genes were assigned to the turquoise module (Fig 4A). Correlations between Module Eigengene, infection and sex were evaluated through Spearman’s correlation analysis (Fig 4B). Infection was significantly correlated with four modules: the turquoise (r= 0.96, p<0.0001), green (r= -0.9, p<0.0001), black (r= 0.71, p<0.0001) and green-yellow (r= -0.68, p= 0.02) (Fig 4B&C). Similar to DEG analysis, sex was not correlated with any of the modules (Fig 4B), suggesting that microglia gene expression is altered by *S. epidermidis* infection in a non-sex-dependant fashion.

**Figure 4.**
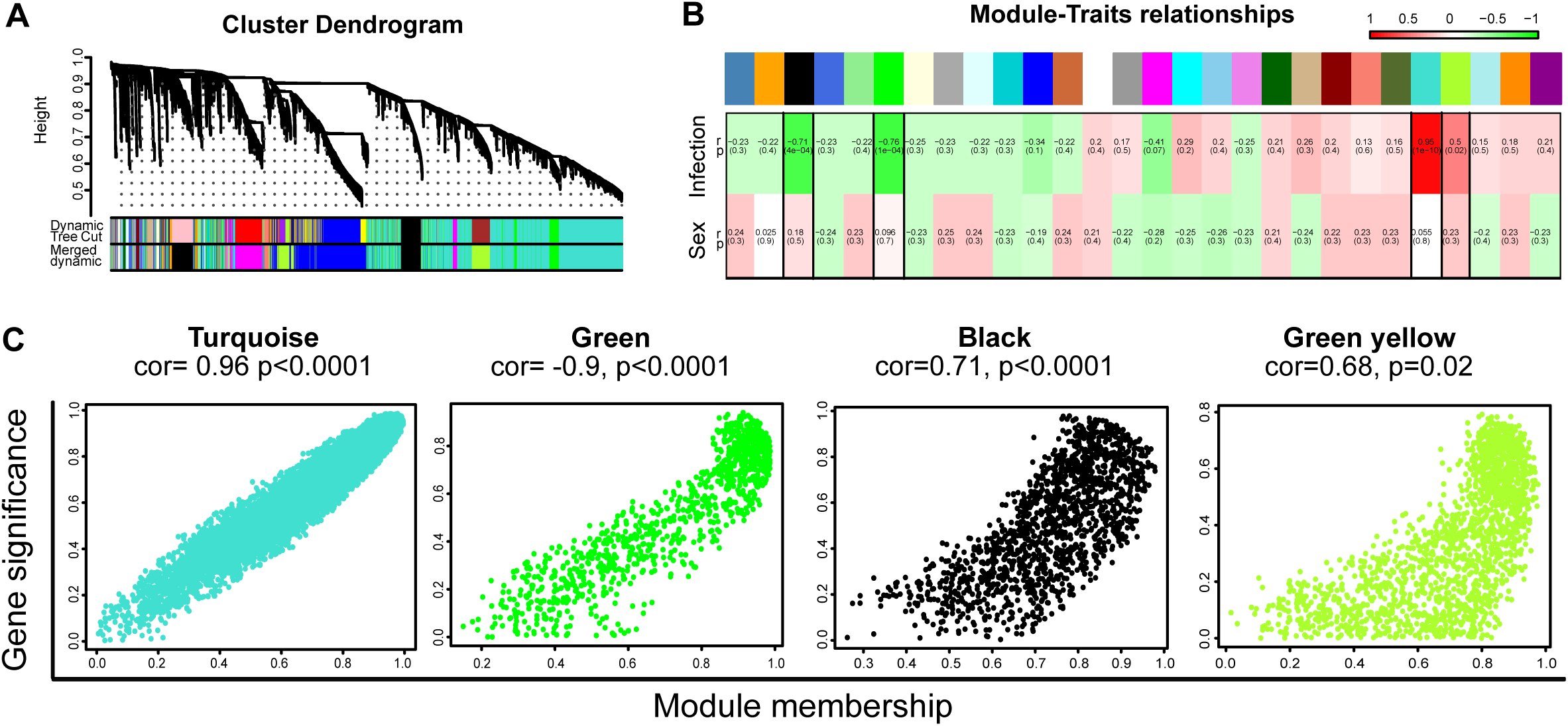
Weighted gene correlation network analysis of hippocampal microglia cells. The cluster dendrogram of the 28 colored modules based on a dissimilarity measure (1-TOM) in microglia cells (A). Correlation between the gene module and clinical traits (Infection and Sex) (B). Scatterplot of gene significance (y-axis) vs. module membership (x-axis) in the most significant modules (Turquoise, green, black and green-yellow) (C).

### 5. Network analysis reveals alteration of crucial pathways following S. epidermidis infection

To evaluate the related molecular mechanisms and pathways, we performed gene ontology (GO) enrichment on the two modules that were most significantly correlated with infection, i.e. turquoise and green. Enrichment map of GO terms in the turquoise module revealed that metabolic processes, protein transport, microglia activation, leukocyte and lymphocyte immunity and cell migration and locomotion were affected after infection (Fig 5A). To identify signalling pathways related to the turquoise module, we annotated the genes by KEGG pathway analysis and found that mechanistic pathways such as Proteasome, NOD-like receptor signalling, RNA transport and spliceosome were altered following *S. epidermidis* infection. Additionally, many neurological disease-related pathways (i.e. Alzheimer disease, Parkinson, spinocerebellar ataxia, Huntington disease) were profoundly affected, suggesting a possible link with neurological deficits (Fig 5B). Enrichment of the green module showed enrichment of genes related to transcription, phosphorylation and biosynthetic processes as well as cell movement/locomotion, axon guidance and cell death (Fig 5C). KEGG pathway analysis of the green module showed alteration in signalling related to development (e.g. Wnt, Notch and axon signalling pathways) and cell adhesion (e.g. Ras, Rap1, focal adhesion pathway, ECM-receptor interaction, tight junction and adherens junction) (Fig 5D). No enrichments were detected in the black and green-yellow modules. Altogether, both WGCNA and DEG analyses indicated involvement of the NOD pathway, leukocytes, cell migration/locomotion and adhesion in activated microglia following *S. epidermidis* infection.

**Figure 5.**
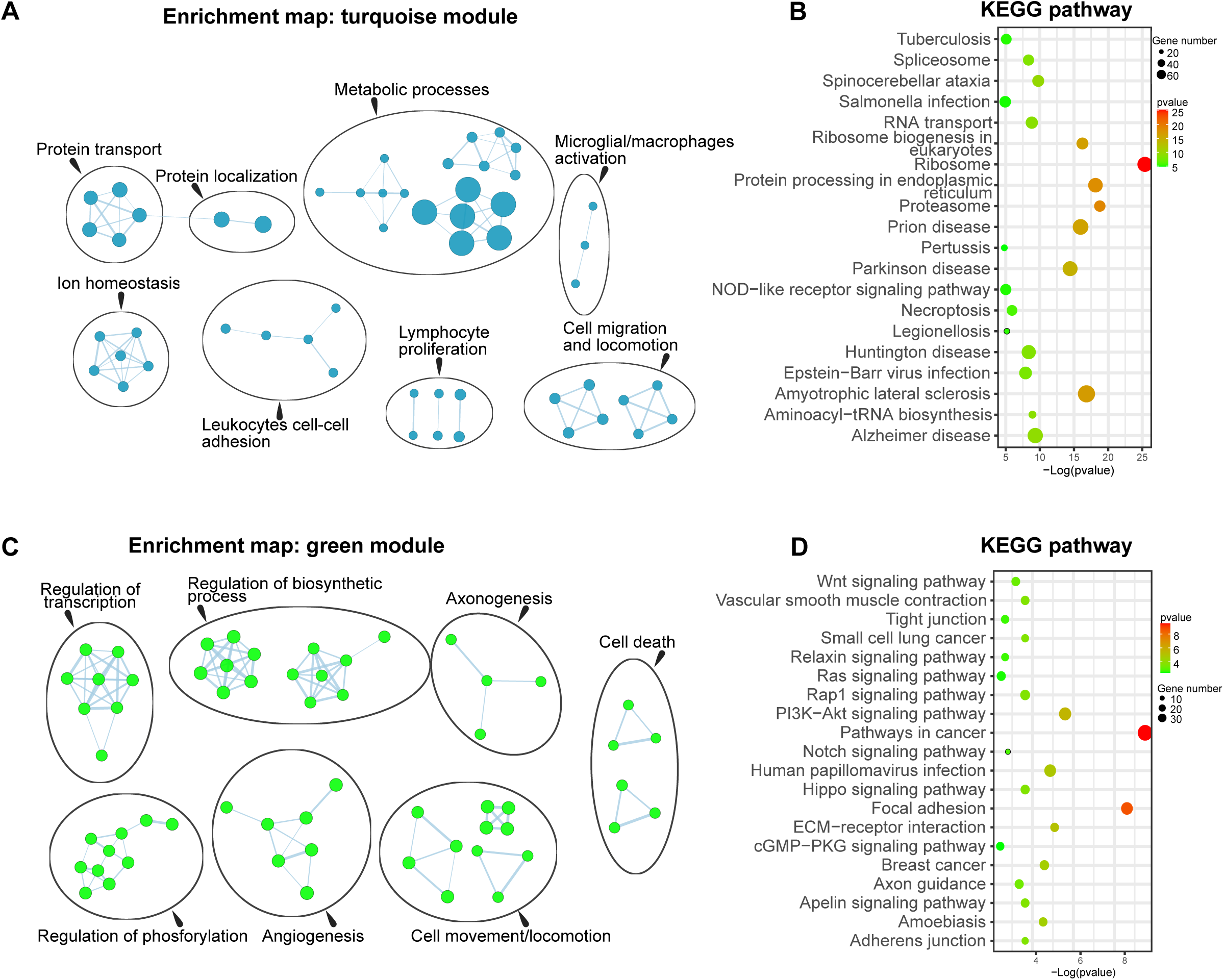
Functional enrichment analysis of most correlated modules. Enrichment map of significantly gene ontology (GO) terms in the turquoise (A) and green (C) modules. Nodes represent gene sets. Highly similar gene sets are connected by edges, grouped in sub-clusters, and annotated automatically. Top 20 significantly enriched KEGG pathways of the turquoise (B) and green (D) modules. Bubble size is proportional to the number of genes in each GO-term, and the color represents the –log(pvalue)

### 6. Peripheral immune cells infiltration into the hippocampus following S. epidermidis infection

To further strengthen DEGs and WGCNA results and gain additional meaningful biological information, we evaluated the degree of overlap between DEGs and the most correlated eigengene modules (Turquoise and Green) (Fig 6A). Both genes in the turquoise and green module exhibited a degree of overlap with DEGs (131 and 58 genes, respectively). Genes in the turquoise module mainly corresponded to upregulated DEGs whilst the genes in the green module mainly overlapped with the downregulated DEGs. To investigate how these genes, interact, we constructed a protein-protein interaction network based on the upregulated-turquoise and downregulated-green shared gene data sets. These genes were highly connected within the respective networks (Fig 6 B & C). To gain further insights into the mechanisms of microglia responses, we also evaluated the degree of overlap between GO biological processes derived from the enrichment analysis of both WGCNA and DEGs analysis. We found that 12 GO biological processes were common to the turquoise and green modules and DEGs. Five were common between the upregulated and downregulated DEGs and the turquoise module, while five between the upregulated, downregulated DEGs and the green module (Fig 6D). Similar to above, there was a striking overrepresentation of biological processes involved in regulation of leukocyte activation, cell proliferation, communication and adhesion, as well as immune system processes, suggesting interaction between microglia and peripheral immune cells.

**Figure 6.**
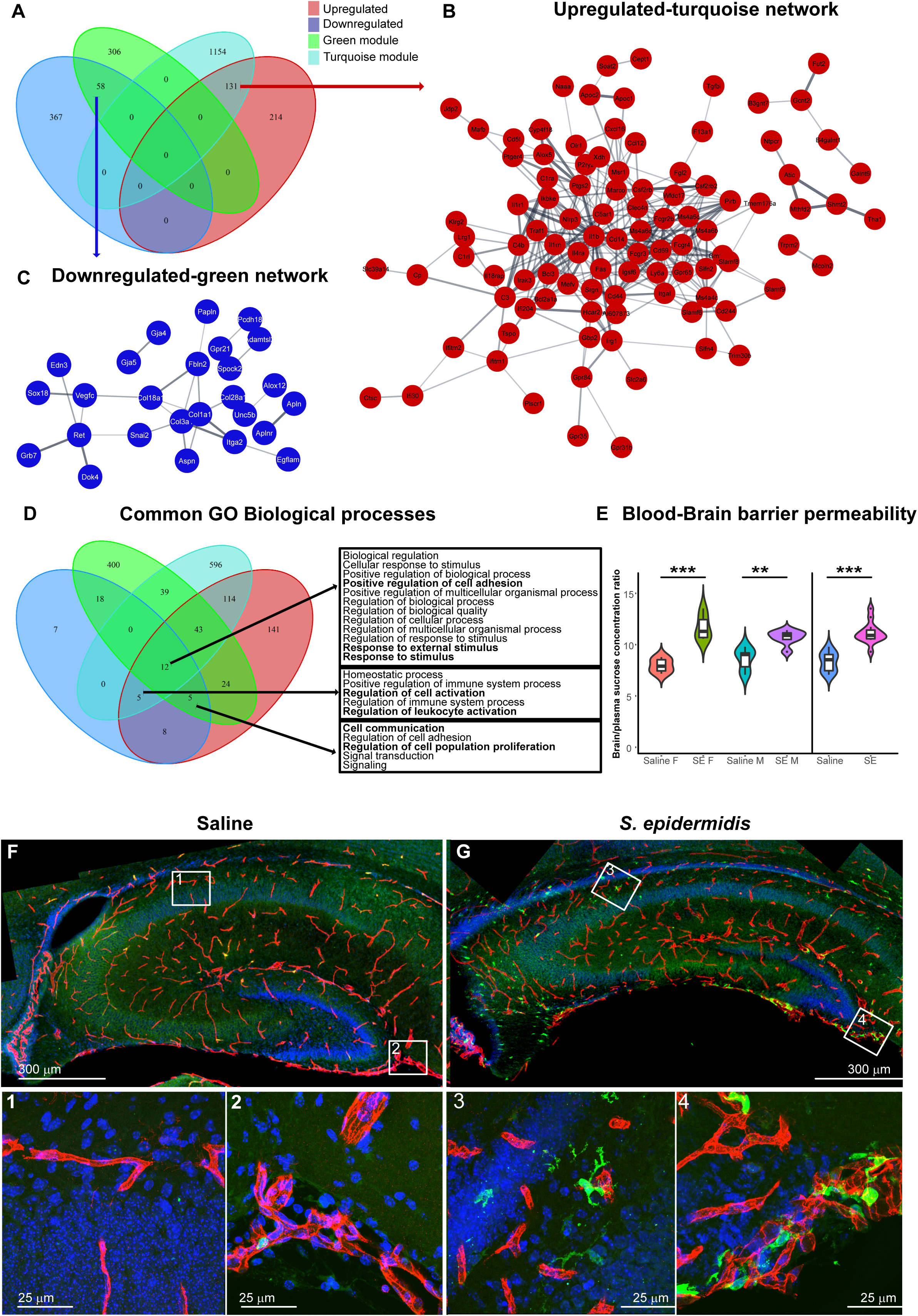
Peripheral immune cells infiltration and blood-brain barrier alteration in the hippocampus in *S. epidermidis* and control mice. Venn diagram demonstrating overlap of all significantly up and downregulated DEGs, turquoise and green module (A). Protein-protein interaction network constructed from the 131overlapping genes between the turquoise module and the upregulated genes (B). Protein-protein interaction network constructed from the 58 overlapping genes between the green module and the downregulated genes (C). The network was created using Cytoscape software. Venn diagram demonstrating overlap of all significantly GO biological processes deriving from the enrichment analysis of the up and downregulated DEGs, turquoise and green module (D). The sucrose brain/plasma concentration ratios in the hippocampus at 24 h after *S. epidermidis* infection (E). Data are presented as median, 10–90th percentile, kernel probability density (violin). Whole-hippocampus immunostaining with anti-CD31 (vessels-Red), anti-GFP (leukocytes-green) and DAPI (nuclei-blue) antibodies shows immune cells infiltration in the hippocampus of *S. epidermidis* infected mice (G) compared to the saline (F). Statistical comparison between the *S. epidermidis* and saline groups for each brain region was performed using Two-way ANOVA with Tukey’s multiple comparison post-hoc test; **p < 0.01, and ***p < 0.001. Saline F: Saline Female; SE F: *S. epidermidis* female; Saline M: Saline male; SE M: *S. epidermidis* male.

Infiltration of leukocytes is an event often observed after blood-brain barrier (BBB) dysfunction, which is known to be associated with several neurological conditions (Rayasam et al., 2021). Therefore, we evaluated BBB permeability after *S. epidermidis* infection by measuring the plasma/brain ratio of glucose in the hippocampus of *S. epidermidis*-infected mice (Ek et al., 2015). Two-way ANOVA analysis revealed a significant main effect of bacterial infection on hippocampal BBB permeability (F value= 53.18, p<0.0001), with increased permeability in infected male and female mice (p=0.003 and p<0.0001 respectively) (Fig 6E).

To further validate the overrepresented biological processes identified by gene analysis, we performed physiological experiments in the LysM-eGFP knock-in transgenic mouse to assess infiltration of peripheral myeloid cells to the hippocampus. In these mice, eGFP is inserted into the lysozyme M locus and labels hematogenous macrophages and neutrophils but not microglia (Faust et al., 2000). Using CD31 immunofluorescence, we distinguished GFP positive leukocytes present in the brain parenchyma from those localized inside the vessels. In hippocampi of saline treated pups, we found single GFP-positive cells mostly localized intravascularly (Fig 6F), while in mice injected with *S. epidermidis*, GFP positive cells were frequently observed in the hippocampus (Fig 6G). All together, these results demonstrate disruption of BBB and that peripheral leukocytes infiltrate the brain parenchyma after *S. epidermidis* infection contributing to the neuroinflammation.

### 7. NOD-signalling pathway activation in microglia of S. epidermidis infected mice

To gain further insight into molecular mechanisms underlying the microglia activation, we evaluated the degree of overlap between KEGG pathways derived from the enrichment analysis using both WGCNA and DEGs analysis. We found 2 pathways in common between the green module and the downregulated network (ECM-receptor interaction, Protein digestion and absorption) and 10 pathways between the turquoise module and the upregulated network, including proinflammatory NF-kappa B, TNF and NOD-like receptor (NLR) signalling pathways (Fig 7A). As NOD signalling has recently been implicated in neonatal infection (Chen et al., 2019) and we found Nlrp3 to be highly connected to the upregulated gene network analysis (Fig 6B), we selected to further investigate genes involved in the NOD pathway. We found that 29/31 genes in the NLR pathway were present in the WGCNA turquoise module, while 10/31 were present in the DEG analysis, and 8 genes were common for both analysis (Fig 7B). Directionality analyses showed that genes in the NOD-signalling pathway were mainly upregulated following *S. epidermidis* infection (Fig 7C). Furthermore, we confirmed that genes in this pathway were highly correlated to each other (Fig 7D), suggesting a clear dependence of these genes within the pathway. Caspase-1 activation is a direct consequence of NLR regulation, (Heneka et al., 2018). Thus, to further validate the involvement of the NOD pathway in microglia activation, we assessed caspase-1 protein concentration in the hippocampus of saline and infected mice. Two-way ANOVA analysis of hippocampal caspase-1 concentration revealed a significant main effect of bacterial infection (F value= 19.411, p<=0.0005). Specifically, at 24 hours after infection, caspase-1 levels were increased in infected male (p=0.01) but not in female mice (p=0.057) (Fig 7E).

**Figure 7.**
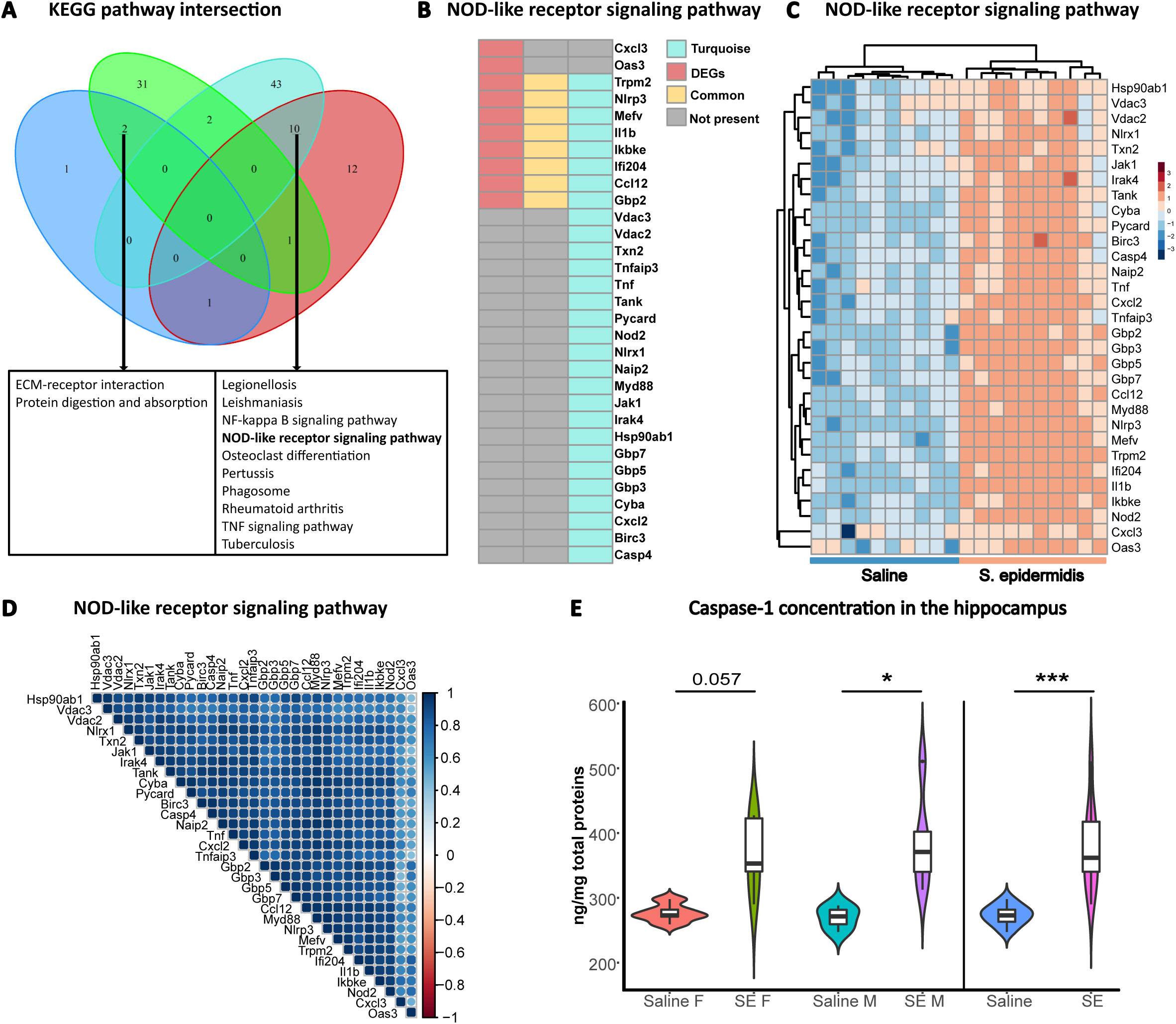
NOD-like receptor signalling activation following S. epidermidis infection. Venn diagram demonstrating overlap of all significantly KEGG pathways from the up and downregulated DEGs, turquoise and green WGCNA module (A). Heatmap showing the contribution of each analysis in the NOD-like receptor pathway (B). Heatmap with directionality (C) and the correlation of the genes involved in the pathway (D). Caspase-1 hippocampal concentration at 24 h after *S. epidermidis* infection (E). Data are presented as median, 10–90th percentile, kernel probability density (violin). Statistical comparison between the *S. epidermidis* and saline groups for each brain region was performed using Two-way ANOVA with Tukey’s multiple comparison post-hoc test; *p < 0.05, and ***p < 0.001. Saline F: Saline Female; SE F: *S. epidermidis* female; Saline M: Saline male; SE M: *S. epidermidis* male.

## Discussion

*S. epidermidis* is a commensal microorganism on the human skin and the most common infectious condition in preterm infants. Infectious disease in preterm infants is a major risk factor for long-term neurological problems, but the underlying mechanisms are not well understood. Here, we deciphered the molecular signature of microglia, the endogenous immune cell in the brain, and discovered that *S. epidermidis* infection profoundly affects the inflammatory response in the developing brain and may play a previously unrecognized important role in neurodevelopmental diseases observed in preterm born children. By combining DEG and WGCNA analyses, we revealed transcriptome reprogramming of microglia and identified inflammasome activation in the hippocampus as a key pathway involved in neonatal *S. epidermidis* infection. Further, we demonstrated morphological evidence of microglia activation in synergy with physiological data on increased BBB permeability and leukocyte infiltration to the brain.

Neuroinflammation in the immature brain is known to affect critical phases of brain development such as myelinisation and neuronal plasticity, as well as induce brain lesions (Hagberg et al., 2015). Epidemiological studies have pointed out infection as a crucial risk factor for neurological conditions in preterm infants such as cerebral palsy (Mitha et al., 2013) and ADHD (Rand et al., 2016), as well as being associated with lower intelligence quotients, memory and attention impairments (Rand et al., 2016, van der Ree et al., 2011). Similarly, prenatal and postnatal exposure to infection has been linked to an increased risk of schizophrenia and autism in mouse and human offspring (Brown and Derkits, 2010, Garbett et al., 2012, Matcovitch-Natan et al., 2016). While *S. epidermidis* is the most common infection in preterm infants, there is scant knowledge of effects on the developing brain and virtually no evidence of pathophysiological mechanisms (Strunk et al., 2014). Several preclinical studies showed that administration of specific toll-like receptor agonists, mimicking gram-positive and negative infections, increas the vulnerability of the brain in neonatal mice (Mottahedin et al., 2017b, Falck et al., 2018, Eklind et al., 2005), rats (Serdar et al., 2019, Eklind et al., 2001) and piglets (Martinello et al., 2019). *S. epidermidis* is believed to act through TLRs and in our seminal studies we showed that this blood borne infection, without crossing the blood-brain barrier, resulted in increased cerebral cytokine and chemokine production and sensitized the brain to hypoxic-ischemic injury (Lai et al., 2020, Gravina et al., 2020, Bi et al., 2015).

Microglia is a key player in neuroinflammation and injury in the immature brain (Van Steenwinckel et al., 2019), however, the effect of *S. epidermidis* infection on microglia is unknown. Microglial dysfunction during development may have marked consequences on developmental programs, which lead to misplaced expression of inflammatory gene pathways and disruption of neuronal development (Kettenmann et al., 2013). Further, as microglia actively communicate with neurons on multiple levels, perturbation of the interplay between microglia and synapses have consequences on aging, neurodevelopmental and neurodegenerative conditions (Cserep et al., 2021). We found that the *S. epidermidis* microglia signature showed similarities with pathways involved in neurological disorders such as Parkinson, Alzheimer, Huntington disease. Hence, the microglia gene signature observed in the present study suggests that *S. epidermidis* may not only have neurodevelopmental consequences in preterm infants, but also longer term effects.

Sex-dependent microglia microRNA differences have been reported to play a role in neurodegenerative diseases (Kodama et al., 2020). Several studies have also reported sex differences in microglia transcriptome in adult animals, although these are not conclusive. Microglia from female mice expressed more proinflammatory genes and immune factors than males (Thion et al., 2018), while others showed that microglia isolated from 12-week-old male mice were more reactive and responsive to stimuli than females (Villa et al., 2018, Guneykaya et al., 2018). In the current study, we did not find sex-dependent differences in the transcriptional program of microglia cells following *S. epidermidis* infection. Neither were there conclusive sex-dependent differences in microglia morphology. Similar to our results, Thion et al. and Hanamsagar et al. showed that during development, microglia do not display transcriptomic differences between sexes and only from postnatal day 60 microglia show differences in their gene expression between males and females (Hanamsagar et al., 2017, Thion et al., 2018).

Microglia are NLRP3 inflammasome-competent cells that exhibit robust canonical NLRP3 inflammasome priming and activation (Hoyle et al., 2020). NLRP3 inflammasome formation is primed by TLR activation (Voet et al., 2019) and subsequently upon inflammasome activation, the adaptor protein ASC initiates formation of active caspase-1, which cleaves pro- IL-1β and pro-IL-18 into bioactive pro-inflammatory cytokines (Heneka et al., 2018). Inflammasome activation, including IL-1β and IL-18, has been implicated in developmental brain injury. Neonatal mice exposed to IL-1β develop long-lasting myelination defects (Favrais et al., 2011). Inflammasome activation was associated with microglia in the injured brain regions in post-mortem tissue of infants with white matter injury and blocking the inflammasome rescued myelination deficit in a mouse model (Holloway et al., 2021). Several studies have also demonstrated beneficial effects on neonatal hypoxic-ischemic brain injury by treatment with the IL-1 receptor antagonist (IL-1ra) (Hagberg et al., 1996), pharmacological inhibition of the NLRP3 inflammasome (Lv et al., 2020) and in transgenic mice deficient in IL-18 (Hedtjarn et al., 2002). In the current study, the transcriptome signature of hippocampal microglia showed extensive involvement of NLR signalling following infection. Accordingly, together with increased levels of caspase-1 protein, we observed upregulation of IL-1 β, Nlrp3 and Pycard genes in microglia cells after *S. epidermidis* infection. Furthermore, inflammasome activation may be enhanced by increased microglia energy metabolism (Ghosh et al., 2018), which we found to be profoundly affected following *S. epidermidis* infection. In support, pharmacological inhibition of energy metabolism derivate has been shown to inhibit NLRP3 inflammasome in microglia (Hoyle et al., 2022). These results suggest that microglia inflammasome activation to be an important mechanism that modulates the neuroinflammatory response following *S. epidermidis* infection.

A fundamental question is to understand is how a blood-borne infection that does not cross the BBB, such as *S. epidermidis*, activates microglia. Here we provide novel microglia transcriptome data that reveals gene networks and pathways involved in regulation of leukocytes and adhesions proteins, suggesting interaction between microglia and peripheral immune cells following *S. epidermidis* infection. In support, using LysM-eGFP mice, we found GFP positive myeloid cells in the hippocampus after infection. While leukocyte trafficking has been reported in several animal models of cerebral ischemia and inflammation in both neonatal and adult mice, contributing to acute and long-term brain injury, underlying mechanisms remain unclear (Rayasam et al., 2021). We previously showed monocyte and granulocyte infiltration to the brain 24 hours after cerebral hypoxia-ischemia (Smith et al., 2018). Following LPS administration to adult animals, Kim et al demonstrated trans-endothelial migration of neutrophils, which interacted with microglia cells and then migrated back to the bloodstream (Kim et al., 2020), possibly spreading inflammatory signals from the brain to the periphery. However, following administration of the TLR1/2 agonist Pam3CSK4 to neonatal mice, peripheral immune cells remained tightly associated with CD31+ brain blood vessels (Mottahedin et al., 2017a), suggesting that *S. epidermidis* infection has broader effects on immune cell trafficking to the brain than activation of specific innate immune receptors. Following *S. epidermidis* infection, we previously showed increased levels of the potent monocyte chemoattractant CCL2 in brain (Gravina et al., 2020), which may have contributed to the leukocyte trafficking in the current study. Further, as CCL2 has been implicated in BBB disruption (Guo et al., 2020), it could also have played a role in the increased permeability of the BBB that we observed, which in turn could facilitate the leukocyte trans-endothelial migration. Interestingly, pharmacological targeting of NLRP3 and caspase-1 prevented microglial activation, reduced infiltration of leukocytes and improved neurologic outcome after cardiac arrest in rats (Chang et al., 2020). Therefore, microglia inflammasome activation, BBB disruption and cell infiltration might be previously unrecognized hallmarks of the complex pathogenesis of *S. epidermidis*-induced neuroinflammation.

Intriguingly, we found marked similarities between *S. epidermidis* and S. aureus (the more virulent cousin) pathways. S. aureus can cause meningitis (Pedersen et al., 2006) and is associated with increased morbidity and mortality in preterm infants (Carey et al., 2008). Activation of microglia cells was observed following S. aureus infection which is able to phagocyte the bacteria and releasing cytokines (Kamata et al., 1985). The immune reaction against S. aureus often includes complement activation a key inflammatory response. In support, we previously showed that complement protein 5 was affected following *S. epidermidis* infection (Gravina et al., 2020). Similarly, complement and coagulation cascade was also an enriched pathway in microglia cells in the current study. Therefore, transcriptome similarities with S. aureus suggests that *S. epidermidis* might be more virulent in the newborn than previously estimated.

In conclusion, we show neuroinflammation following exposure to the commensal microorganism *S. epidermidis*, which is the most common infectious condition in preterm infants. Our study identify microglia inflammasome activation in association with increased BBB permeability and trans-endothelial trafficking of myeloid cells into the brain parenchyma as key mechanisms that underlie neuroinflammation following *S. epidermidis* infection. The results demonstrate that neonatal *S. epidermidis* infection share many transcriptome pathways with S. aureus and several neurodegenerative diseases, suggesting a previously unrecognized important role in neurodevelopmental disorders observed in preterm born children.

## Material and methods

### Animals

C57Bl/6J wild-type mice were purchased from Janvier Labs (Le Genest-Saint-Isle, France) and Charles River Laboratories (Sulzfeld, Germany) and were bred at the animal facility at the University of Gothenburg (Experimental Biomedicine, University of Gothenburg). LysM-EGFP-ki mice were purchased from University of Missouri Mutant Mouse Regional Resource Center and bred internally. Mice were housed with a normal 12-h light/dark cycle (lights on at 06:00) and ad libidum access to standard laboratory chow diet (B&K, Solna, Sweden) and drinking water in a temperature-controlled environment (20–22°C). All animal experiments were approved by the Gothenburg Animal Ethical Committee (No 663/2017). Mice of both sexes were used. Sex was established by visual inspection. In each experimental group, mice were obtained from at least three different litters.

### Bacterial growth

For exposure to neonatal infection, mice were intraperitoneally injected with sterile saline or 3.5 × 10^7^ colony-forming units (CFU) of *S. epidermidis* at postnatal day (PND) 4 as previously described (Lai et al., 2020).

### Sample collection for histological evaluation

At PND5, C57Bl/6J wild-type and LysM-EGFP-ki pups were deeply anesthetized via intraperitoneal administration of pentobarbital (Pentacour) and perfused transcardially using saline. Brains were collected, post-fixed in 6% buffered formaldehyde (Histofix; Histolab products AB, Västra Frölunda, Sweden) followed by immersion in 30% sucrose for 48 hours for cryoprotection frizzing in cold isopentane (Sigma Aldrich, Darmstadt, Germany). Using cryostat (Leica, CM 3050 S, Germany), 40-µm thick coronal sections were obtained. The first section of each series was randomly chosen by using a random table and the section sampling fraction (SSF) was one out of four.

### Immunofluorescence

One set of sections from LysM-EGFP-ki pups was used to detect presence of GFP-positive cells in the hippocampus. Selected brain sections were permeabilized in citrate buffer at 85°C, blocked in 4% normal donkey serum, incubated in 0.2 % Triton X-100 with goat anti-CD31 (1:250; AF3628; R&D systems) overnight at 4°C followed by incubation with donkey anti-goat Alexa Fluor 594 secondary antibodies (1:200, A-11058; Invitrogen) and rabbit anti GFP (1:100, A-21311; Invitrogen) antibodies, counterstaining with DAPI solution and mounting on slides using Antifade Mounting Medium (Vector Laboratories). Brain sections were examined and visualized using laser scanning confocal microscope Zeiss (LSM 800). Z-stacks of images were captured from brain sections with z-plane step size of 4.0 µm using 10× air objective lens, or 1.0 µm using 40× oil-immersion objective lens. Images were processed using Fiji software version 1.53c and maximum projection of z-stacks were made to present the results (Schindelin et al., 2012). Large field images covering the whole hippocampus area were created from low magnification images using Fiji stitching plug-in (Preibisch et al., 2009).

### Immunohistochemistry

One set of sections from C57Bl/6J wild-type mice was stained for ionized calcium binding adaptor molecule 1 (Iba1). The sections were rinsed in PBS (2 × 10 min) before antigen-retrieval by incubation in target retrieval solution (Dako S1700, Glostrup, Denmark) for 40 min at 85 °C. Slides were then rinsed (2 × 10 min in PBS) followed by a 10 min block with endogenous peroxidase (1 mL 30% H_2_O_2_ and 9 mL PBS), a rinse in washing buffer (0.25% T-PBS for 20 min). Afterwards, sections were incubated overnight in the rabbit anti-Iba1 polyclonal antibody as a primary antibody (diluted 1:500 in washing buffer) (ab108539, Abcam). The next day, sections were rinsed with washing buffer (2 × 10 min) and incubated in goat anti-rabbit IgG (1:200) for 2 h at room temperature. Subsequently, sections were rinsed in washing buffer (2 × 10 min) followed by addition of ABC solution (VECTASTAIN Elite ABC HRP Kit (Peroxidase, Standard, PK-6100) for 1 h and rinsing in washing buffer for 20 min. In the last step, immunolabelling using 3,3′-diaminobenzidine (DAB) solution (DAB EASY tablets, ACROS Organics™) for 2 minutes was performed. Finally, the sections were mounted on the gelatin-coated slides, dried for 20 min, re-hydrated in demineralized water for 2 min, counter stained with 0.25% thionin solution (thionin, Sigma T3387), dehydrated through a graded series of alcohol (96%, 99%), cleared in xylene for 10 min and coverslipped

### Measurement of the size of microglia

The quantification of the size of microglia soma was performed on Iba-1 stained sections by measuring the volume of the cell soma with a 3D nucleator method, in two subregions of hippocampus, the CA1 stratum radiatum (CA1.SR) and the molecular layer of dentate gyrus (MDG). Delineation of CA1.SR and MDG subregions were performed using light microscope under 5 × objective lens, the newCAST software (Visiopharm, Hørsholm, Denmark) modified for stereology with a digital camera (Leica DFC 295, Germany) and a motorized stage (Ludl Mac 5000, US). By using 3D nucleator, the number of half-lines was set at 6 and the mode was vertical uniform random (VUR) based on the assumption of rotational symmetry of microglia. Volume of Iba-1-immunopositive microglia were estimated with a 100 × oil-immersion objective lens. For each animal, 50-80 microglia were randomly sampled by using optical disector.

### 3-D reconstruction of microglia

The morphology of the CA1SR hippocampal microglia was quantified on 10 microglia per animal in terms of the number of branches, total branch length, soma sphericity and the branching complexity. This was done using the Filament Tracers algorithm (supplementary video 1) and Surface module of the Imaris software (Version 8.4, Bitplane A.G., Zurich, Switzerland) on the images captured with set of Z-stacks of images (step size 1 µm) using a ×63 oil-immersed lens. The complexity of microglia processes was quantified using Sholl analysis (Gober et al., 2022). To investigate microglia sphericity, three-dimensional ellipsoid plots were generated by MATLAB (R2020b) according to: https://mathworld.wolfram.com by using collected information from Imaris ellipsoid function.

### Hippocampi microglia isolation

Hippocampi from saline-perfused C57Bl/6J wild-type mice were extracted and placed in HBSS without Ca and Mg and immediately processed for tissue dissociation and CD11b+ isolation. Hippocampi tissues were dissociated to single-cell suspension by enzymatic degradation using a MACS Technology neural tissue dissociation kit (Miltenyi Biotec, Cat# 130-092-628) according to the manufacturer’s protocol. Pre-warmed enzyme mix was added to the tissue pieces, which were incubated with agitation at 37°C. The tissue was further mechanically dissociated by trituration, and the suspension was applied to a 40-μm cell strainer placed on a 50 mL tube. 10 mL HBSS (with Ca and Mg) was added through the strainer and centrifuged for 10 min at 300g at room temperature. Before cell isolation, cells were counted by mixing 5 µL cell suspension with 5 µL Tryptan Blue, the mixture was added to BioRad counting slides and counted in a BioRad cell counter (Bio-Rad Laboratories, Hercules, Ca). Following counting, cells were immediately processed for MACS MicroBead separations. To separate primary microglia, the CD11b+ cells were magnetically labelled with CD11b (microglia) MicroBeads. The cell suspension was loaded onto a MACS Column (MS column) (Miltenyi Biotec), which was placed in the magnetic field of a MACS Separator (Miltenyi Biotec). The magnetically labelled CD11b+ cells were retained within the column. After removing the column from the magnetic field, the magnetically retained CD11b+ cells were eluted as the positively selected cell fraction. CD11b+ cells were counted in BioRad cell counter as described above. The CD11b+ cell pellets were then stored at −80°C for later RNA extraction.

### RNA extraction and RNA sequencing

Isolated CD11b+ cell pellets were homogenized in RLT Lysis buffer. Two tubes with CD11b+ cell lysed pellets were pooled in the same column for RNA extraction. Total RNA was purified by miRNeasy Micro Kit according to the manufacturer’s protocol. RNA quantity was analyzed by qubit (Invitrogen). Preparation of RNA library and transcriptome sequencing was conducted by Novogene Co., LTD (Cambridge, UK). The library prep was stranded and sequencing depth was 60M pair-end reads (2*250).

### Read alignment and expression quantification

The raw data were saved as FASTQ format files and FastQC (version 0.11.2) were used for quality control. Trimming of the adapter content and Quality trimming was performed using TrimGalore (version 0.4.0) tool, a wrapper around Cutadapt (version 1.9) (Martin, 2011). Trimmed reads were mapped to the Mus-musculus mm10 genome using STAR (version 2.5.2b) (Dobin et al., 2013). FeatureCounts (version 1.6.4) (Liao et al., 2014) was used to quantify number of reads mapped to each gene using a list of annotated genes downloaded from Ensembl (version 83) Mus-musculus GRCm38 (Flicek et al., 2014).

### Differential gene expression analysis

The overall dataset contained 46984 probe sets. In order to focus on annotated functional genes, the dataset was filtered using the STRING app on cytoscape software. After filtering, we identified 20834 transcripts, which was used for further exploration. Out of 20834, 17643 with nonzero total read count were used for the analysis. Differential expression were generated using DESeq2 v1.32.0 from Bioconductor (Love et al., 2014). For pairwise comparisons, we compared *S. epidermidis* infected mice to saline mice in males and females (padj < 0.05, FC > 2 in both comparisons). Data were visualized using “pheatmap” and “EnhancedVolcano” packages.

### Weighted gene co-expression network analysis (WGCNA)

Standard WGCNA procedure was followed to create signed gene co-expression networks from the WGCNA R package v1.68 (Langfelder and Horvath, 2008). A total of 20834 transcripts were imported for WGCNA analysis. Using the function “goodSamplesGenes”, 3190 transcripts were excluded due to zero variance, therefore 17644 transcripts were used for the analysis. The basic principle of WGCNA is to build a network of genes where values derived from correlations of expression profiles are used as edges. The correlation/similarity values are however transformed in two steps, into adjacency and topological overlap respectively, to ensure that the network has what is considered natural/expected properties. Transformation to adjacency involves raising similarity to a power β that should be chosen so that the derived network has a scale free topology. WGCNA provides a function that generates a plot to aide in selecting a proper value for β. In this analysis β was set to 4. The resulting signed adjacency matrix was used to calculate topological overlap matrix (TO), which was then hierarchically clustered with (1-TO) as a distance measure. Subclusters or modules were derived with a tree cutting algorithm. A representative gene expression profile, or eigengene, was then derived for each module as the first principal component of the gene expression values within the module. Pearson correlation between each gene and module eigengene was calculated to identify module membership, which represents how close a particular gene is to a module. Due to the high number of genes in each module, genes with module membership higher than 0.9 were selected for enrichment analysis.

### GO enrichment

Gene ontology (GO) and Kyoto Encyclopedia of Genes and Genomes (KEGG) pathway enrichment analyses were performed using Cytoscape StringApp (Doncheva et al., 2019) and visualized in R software. GO terms and KEGG pathways with P. adjust to < 0.05 were considered to be significantly enriched. To overcome gene-set redundancy and help in the interpretation of large gene lists, Enrichment Map was used to visualize GO biological processes (Merico et al., 2010). Cyto-Hubba plug-in of the software Cytoscape was used to screen for hub genes in the upregulated and downregulated network based on the Maximum Neighborhood Component (MNC) algorithm. Furthermore, to limit the study to common processes, GO terms and KEGG pathways with P. adjust to < 0.05 from both DEGs and WGCNA were merged and visualized using “VennDiagram” package.

### Blood-brain barrier permeability

A quantitative assessment of regional BBB function was performed by measuring sucrose blood to hippocampus uptake at 24 h after *S. epidermidis* infection or saline injected control animals. Briefly, animals were injected with ^14^C-sucrose (342 Da) intraperitoneally (0.2μCi/g body weight; 10μL injection/g animal) and killed 30 minutes later by pentobarbital injection. Blood was collected from the heart through cardiac puncture and spun to separate plasma. The brains were dissected out and the hippocampus was to assess regional BBB permeability. Brain tissue samples were dissolved using Solvable (0.5mL; PerkinElmer) and 4.5mL scintillation fluid added to all samples (UltimaGold, PerkinElmer). Isotope activity in each fluid or tissue sample was determined by liquid scintillation counting (LSC) and brain/plasma sucrose concentration ratios were used as a measure of BBB permeability as previously described (Ek et al., 2015)

### Measurement of hippocampal Caspase-1

Caspase-1 activity from hippocampal lysates was analysed by enzyme-linked immunosorbent assay (ELISA) kit and standard (Mouse caspase-1, Novus Biologicals, Littleton, CO, USA, catalog # NBP2-75014) as per manufacturer’s instructions.

### Statistical analyses

Statistical analyses were performed in R software (version 1.4.1106). For the microglia morphology, BBB permeability and caspase-1 concentration, Two-way ANOVA with Turkey post-hoc test was performed using the functions “aov” and “TukeyHSD”. Data are presented as median, 10–90th percentile, kernel probability density (violin). P < 0.05 was considered statistically significant.

## Acknowledgments

We thank Novogene Co., LTD (Cambridge, UK) for performing the sequencing. We also acknowledge Katarine Tuvé from the Genomics and Bioinformatics Core Facility platforms at the Sahlgrenska Academy, University of Gothenburg, for performing the read alignment and expression quantification of the sequencing raw data. We also acknowledge Dr Ali Rafati for providing microglia sphericity, Kajsa Groustra for help with mouse breeding (Experimental Biomedicine, Sahlgrenska Academy, University of Gothenburg) and Anna-Lena Leverin for technical support. Graphical abstract was created with BioRender.com (Agreement number: MZ23M9V8RZ).

## Author contribution

The study was conceived and designed, and the manuscript drafted by GG and CM. Experiments were performed by GG, MA, TC and CJE. Data analysis was performed by GG, MA, TC, HR and CJE. All authors critically reviewed and edited the work.

## Data and Code Availability

The data that support the findings of this study are available from the authors CM and GG upon request.

---

**Supplementary table 1.**

| **Hub genes in the upregulated network** |  |  |  |
| --- | --- | --- | --- |
| **Gene Name** | **Uniprot description** | **Log2FC** | **Padj** |
| ***Csf3*** | Colony stimulating factor 3 (granulocyte); Granulocyte/macrophage colony-stimulating factors are cytokines that act in hematopoiesis by controlling the production, differentiation, and function of 2 related white cell populations of the blood, the granulocytes and the monocytes-macrophages. This CSF induces granulocytes; Belongs to the IL-6 superfamily. | 3.38 | 1.64E-43 |
| ***ApoB*** | Apolipoprotein B-100; Apolipoprotein B is a major protein constituent of chylomicrons (apo B-48), LDL (apo B-100) and VLDL (apo B-100). Apo B-100 functions as a recognition signal for the cellular binding and internalization of LDL particles by the apoB/E receptor. | 1.01 | 3.86E-11 |
| ***Crp*** | C-reactive protein, pentraxin-related; Displays several functions associated with host defense: it promotes agglutination, bacterial capsular swelling, phagocytosis and complement fixation through its calcium-dependent binding to phosphorylcholine. Can interact with DNA and histones and may scavenge nuclear material released from damaged circulating cells; Belongs to the pentraxin family. | 5.2 | 7.15E-08 |
| ***C3*** | Complement component 3; C3 plays a central role in the activation of the complement system. Its processing by C3 convertase is the central reaction in both classical and alternative complement pathways. After activation C3b can bind covalently, via its reactive thioester, to cell surface carbohydrates or immune aggregates. | 1.24 | 1.50E-21 |
| ***Fcgr2b*** | Low affinity immunoglobulin gamma Fc region receptor II; Receptor for the Fc region of complexed immunoglobulins gamma. Low affinity receptor. Involved in a variety of effector and regulatory functions such as phagocytosis of antigen-antibody complexes from the circulation and modulation of antibody production by B-cells. Isoform IIB1 and isoform IIB1' form caps but fail to mediate endocytosis or phagocytosis. Isoform IIB2 can mediate the endocytosis of soluble immune complexes via clathrin- coated pits. Isoform IIB1 and isoform IIB2 can down-regulate B- cell, T-cell, and mast cell activation when coaggregated to B-cell receptors for AG (BCR), T-cell receptors for AG (TCR), and Fc receptors, respectively. | 1.36 | 2.87E-58 |
| ***Cd14*** | Myeloid cell-specific leucine-rich glycoprotein; Coreceptor for bacterial lipopolysaccharide. In concert with LBP, binds to monomeric lipopolysaccharide and delivers it to the LY96/TLR4 complex, thereby mediating the innate immune response to bacterial lipopolysaccharide (LPS). Acts via MyD88, TIRAP and TRAF6, leading to NF-kappa-B activation, cytokine secretion and the inflammatory response. Acts as a coreceptor for TLR2:TLR6 heterodimer in response to diacylated lipopeptides and for TLR2:TLR1 heterodimer in response to triacylated lipopeptides, these clusters trigger signaling from the cell surface and subsequently are targeted to the Golgi in a lipid-raft dependent pathway (By similarity). Acts as an accessory receptor for M.tuberculosis lipoproteins LprA, LprG and LpqH, in conjunction with coreceptors TLR2 and TLR1. The lipoproteins act as agonists to modulate antigen presenting cell functions in response to the pathogen. Binds electronegative LDL (LDL(-)) and mediates the cytokine release induced by LDL(-) (By similarity). | 1.17 | 6.61E-35 |
| ***Il1b*** | Interleukin 1 beta; Potent proinflammatory cytokine. Initially discovered as the major endogenous pyrogen, induces prostaglandin synthesis, neutrophil influx and activation, T-cell activation and cytokine production, B-cell activation and antibody production, and fibroblast proliferation and collagen production; Belongs to the IL-1 family. | 2.09 | 6.38E-106 |
| ***Fcgr4*** | Low affinity immunoglobulin gamma Fc region receptor IV; Receptor for the Fc region of immunoglobulin gamma. Also acts as a receptor for the Fc region of immunoglobulin epsilon. Binds with intermediate affinity to both IgG2a and IgG2b. Can bind to IgG2a and IgG2b monomers. Does not display binding to IgG1 or IgG3. Mediates neutrophil activation by IgG complexes redundantly with Fcgr3. Plays a role in promoting bone resorption by enhancing osteoclast differentiation following binding to IgG2a. Binds with low affinity to both the a and b allotypes of IgE. Has also been shown to bind to IgE allotype a only but not to allotype b. Binds aggregated IgE but not the monomeric form and bound monomeric IgG is readily displaced by IgE complexes. Binding to IgE promotes macrophage-mediated phagocytosis, antigen presentation to T cells, production of proinflammatory cytokines and the late phase of cutaneous allergic reactions. | 2.12 | 7.30E-136 |
| ***Fcgr3*** | Low affinity immunoglobulin gamma Fc region receptor III; Receptor for the Fc region of complexed immunoglobulins gamma. Low affinity receptor which binds to IgG1, IgG2a and IgG2b. Mediates neutrophil activation by IgG complexes redundantly with Fcgr4. | 1.06 | 4.08E-47 |
| ***Sell*** | Leukocyte-endothelial cell adhesion molecule 1; Cell surface adhesion protein. Mediates the adherence of lymphocytes to endothelial cells of high endothelial venules in peripheral lymph nodes. Promotes initial tethering and rolling of leukocytes in endothelia (By similarity); Belongs to the selectin/LECAM family. | 1.16 | 2.13E-09 |

**Supplementary table 2.**
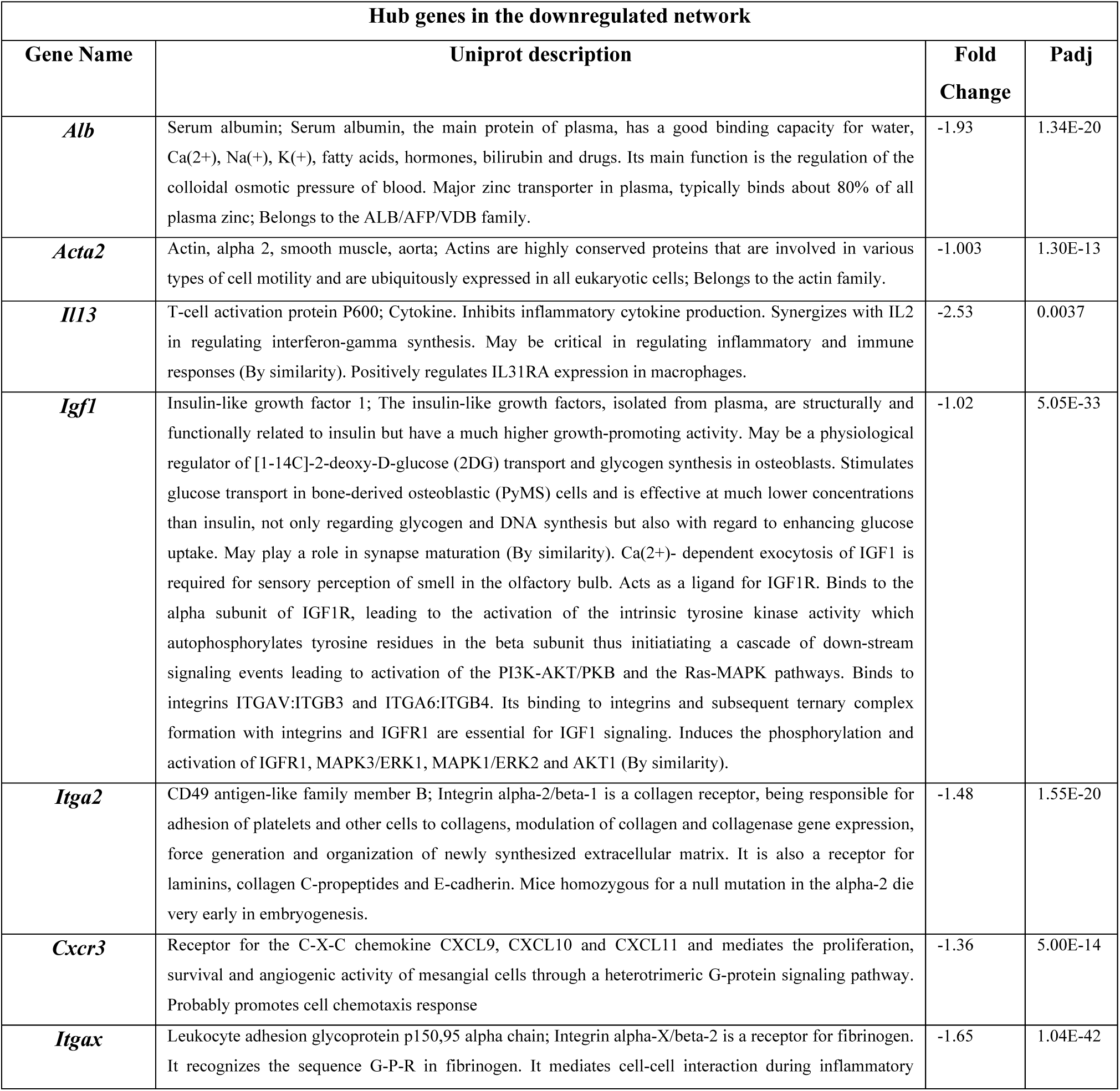
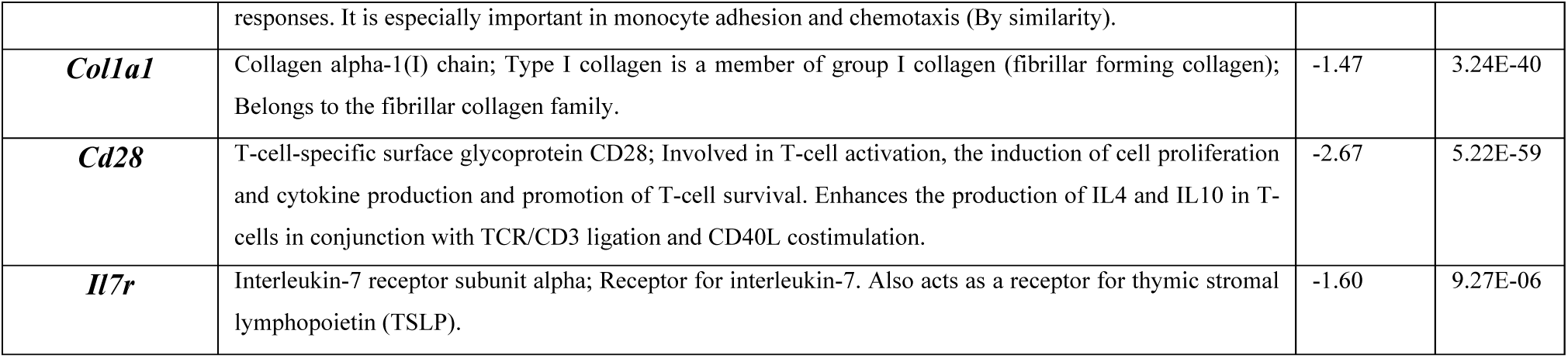

**Figure S1.**
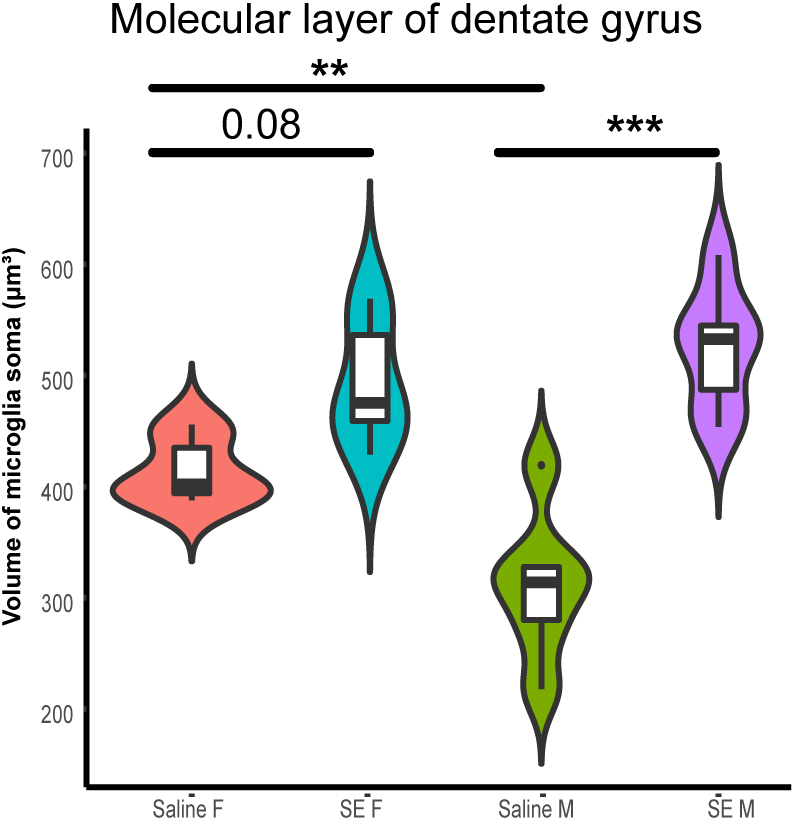
Activation of hippocampal microglia cells following S. epidermidis infection in the molecular layer of dentate gyrus. Microglia volume was quantified in PND5 mice (n=6 sex/treatment) in the molecular layer of dentate gyrus (MDG) hippocampal subregion. Data are presented as median, 10–90th percentile, kernel probability density (violin). Statistical comparison between the *S. epidermidis* and saline groups for each parameter was performed using Two-way ANOVA with Tukey’s multiple comparison post-hoc test. *p<0.05; **p<0.01; ***p<0.001

**Figure S2.**
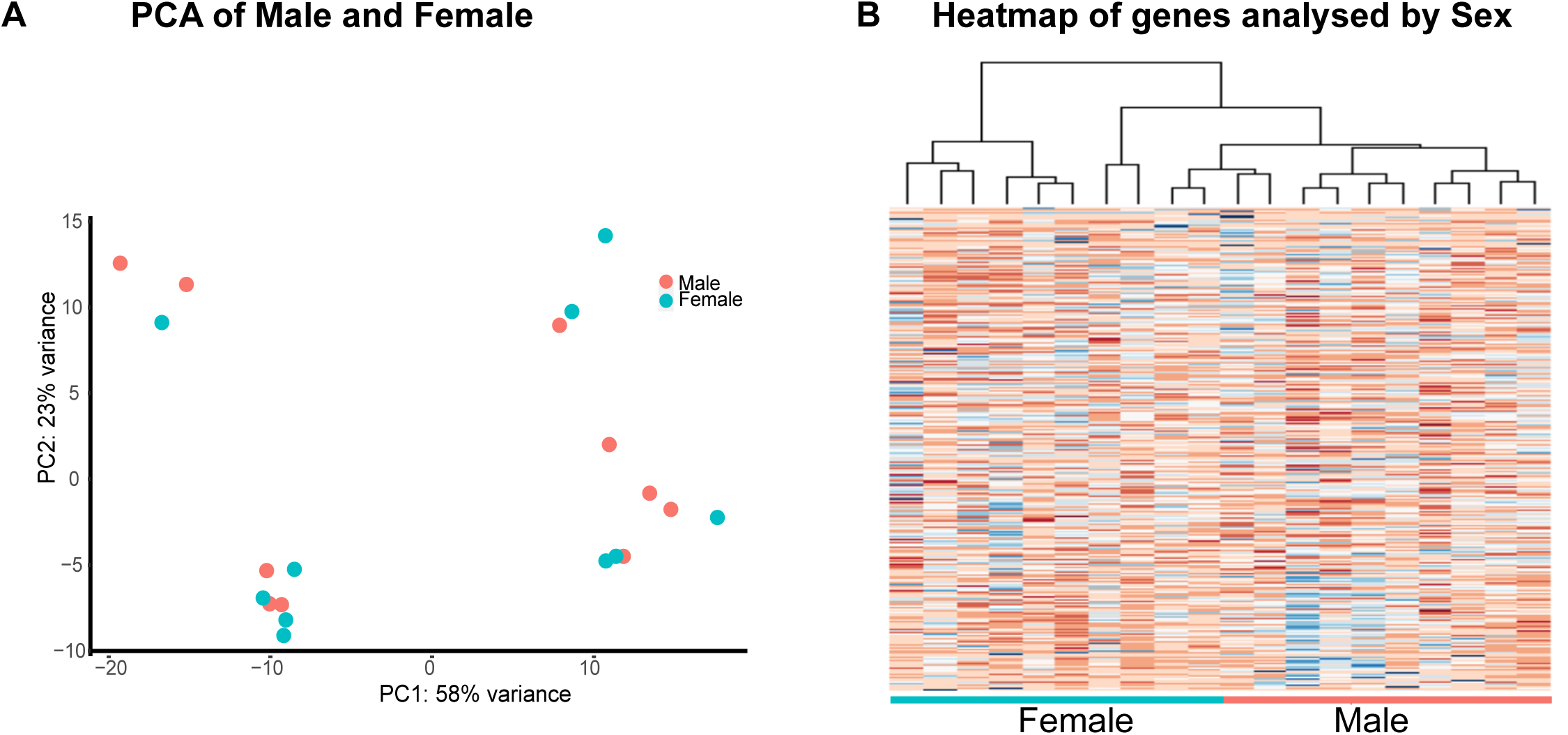
S. epidermidis affects hippocampal microglia transcriptome in a non-sex-dependant fashion. (A) Principal component analysis (PCA) shows no separation of microglia transcripts of male and female mice (saline n = 10, *S. epidermidis* n = 10). B. The heatmap shows also no difference between male and female microglia transcriptome

**Figure S3.**
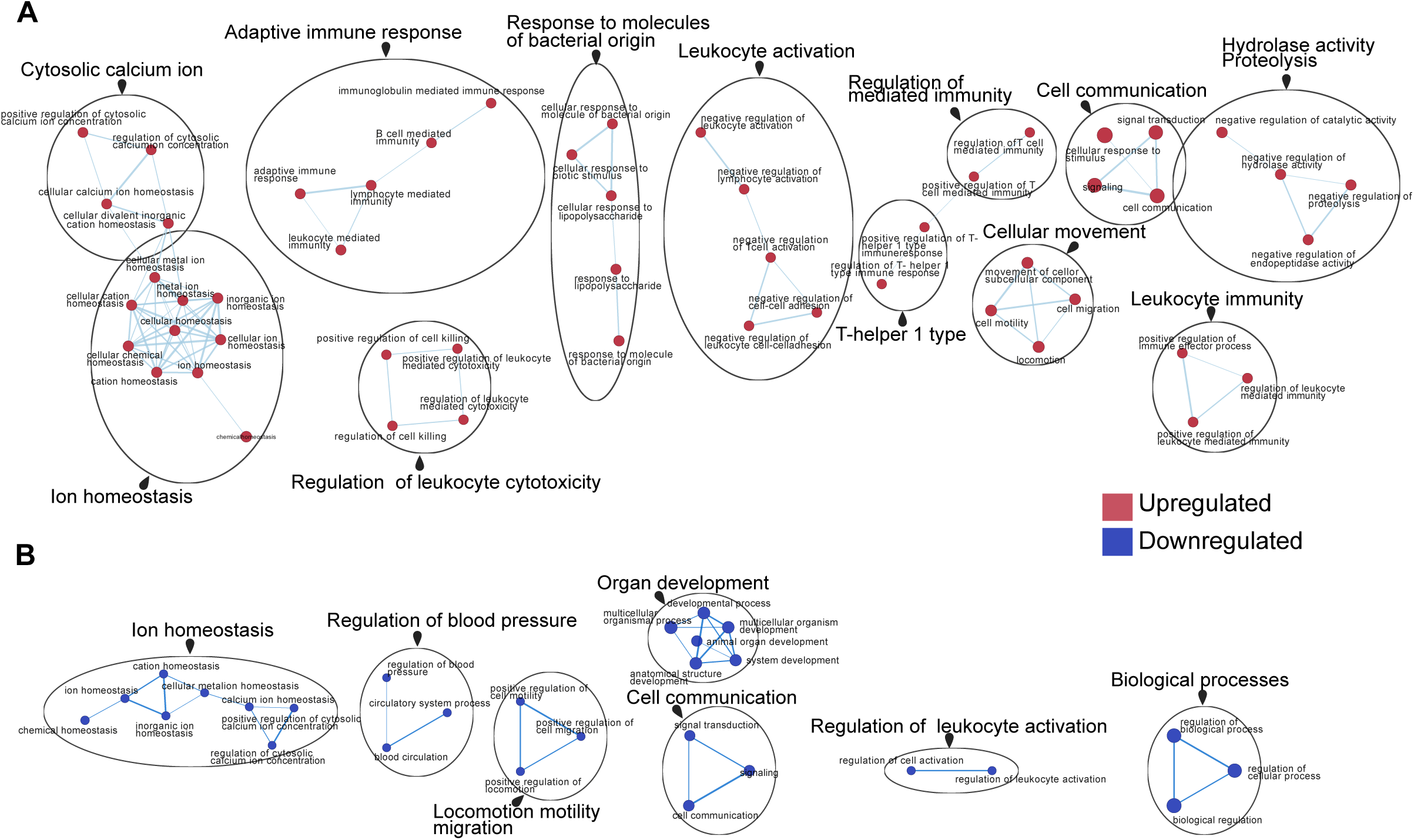
Functional enrichment analysis of differential expressed genes. Enrichment map of significantly gene ontology (GO) terms in the upregulated (A) and downregulated (B) network. Nodes represent gene sets. Highly similar gene sets are connected by edges, grouped in sub-clusters, and annotated automatically.

